# Changes in biochemistry and histochemical characteristics during somatic embryogenesis in *Ormosia henryi* Prain

**DOI:** 10.1101/2020.09.21.307009

**Authors:** Gaoyin Wu, Xiaoli Wei, Xiao Wang, Xian Liang, Yi Wei

**Affiliations:** College of Forestry/Institute for Forest Resources & Environment of Guizhou, Guizhou University, Guiyang 550025, Guizhou, People’ s Republic of China

**Keywords:** *Ormosia henryi*, Somatic embryogenesis, Biochemical events, Histochemical observation

## Abstract

Mature embryos were used as an explant for embryogenic callus (EC) induction, and then EC was further developed to form somatic embryos during somatic embryogenesis (SE) of *Ormosia henryi* Prain; however, some mature embryos could induced non-embryogenic callus (NEC), browning callus (BC) or snowflake callus (SC). These phenomena might be related to the biochemical and histochemical differences during somatic embryo induction. The present study was conducted to analyze the biochemical events and histochemical changes at different SE stages in *0. henryi*. The contents of soluble sugar, starch, soluble protein, H_2_O_2_, and endogenous hormones and the activities of polyphenoloxidase (PPO), superoxide dismutase (SOD), ascorbate peroxidase (APX), peroxidase (POD) and catalase (CAT) were measured at different SE stages, such as EC, globular embryo (GE), and cotyledon embryo (CE), and in abnormal tissue, such as NEC, BC, and SC. The results showed that the contents of soluble sugar and starch; the activities of PPO, SOD, APX and POD; and the ratios of indole-3-acetic acid/abscisic acid (IAA/ABA), IAA/gibberellins (IAA/GAs), auxin /GAs (AUX/GAs), and AUX/ABA decreased gradually at different SE stages. In contrast, the contents of soluble protein, H_2_O_2_, all endogenous hormones gradually increased. However, CAT activity and the ratios of IAA/cytokinins (IAA/CKs), AUX/CKs, ABA/CKs, and GAs/CKs first increased and then decreased. The high contents of GAs and ABA, high ratios of ABA/CKs and GAs/CKs and low ratios of IAA/ABA, IAA/GAs, AUX/GAs and AUX/ABA were responsible for the inability of the callus to form EC. The low enzyme activities, low contents of energy substances and H_2_O_2_ were related to NEC formation. The high contents of soluble sugar, H_2_O_2_, AUX, CKs and PPO activity and the low content of soluble protein were the basic causes of BC formation. The high-energy substances contents and low activities of SOD and POD facilitated SC formation. Histochemical observation showed that starch granule staining gradually lightened with SE development, but protein granules were darkly stained. Compared with EC, starch and protein granules were stained darker in SC, and lighter in NEC and BC. These results showed that energy substances were the material basis of SE, which affected enzyme activities, regulated reactive oxygen species (ROS) metabolism, and thus regulated the morphogenesis and development of somatic embryos. In addition, the contents and ratios of endogenous hormones affected the dedifferentiation, dedifferentiation and embryogenesis of somatic cells. To induce EC from mature embryos and further develop their formation into somatic embryos, it is necessary to adjust the energy supply and hormone ratio in the medium.

**One-sentence summary:** Somatic embryogenesis and abnormal callus tissues formation of *Ormosia henryi* Prain were associated with energy substances, reactive oxygen species, enzyme activities and endogenous hormones, as well as histochemical characteristics.

## INTRODUCTION

Somatic embryogenesis (SE) is an efficient approach for the clonal propagation of plants (Correia et al., 2011), Its developmental process resembles zygotic embryo development (Leljak-Levanic et al., 2004), which is an ideal material for studying morphology, physiology, biochemistry and molecular biology in embryogenesis and development (Morel et al., 2014). Since the late 1950s, Steward (1958) found that carrot root cells could be regenerated through SE in vitro, and the SE for many plants has been studied. *O. henryi* is a precious tree species in China and has great economic and ecological potential as a raw material for making high-grade furniture and handicrafts; some factors restrict the scale propagation of *O. henryi*, such as scarce wild resources, biennial fruiting, dormancy caused by the hardness seed coat and pests that eat the seeds (Wei et al., 2014). Therefore, it is very important to research asexual propagation for *O. henryi*. Our previous research has established the SE protocols of *O. henryi* (Wu et al., 2020), but the underlying mechanism is not well understood. During the process of callus subculture, callus is prone to browning and eventually necrosis, and its transformation into SC is a common phenomenon, seriously hindering the further development of callus. Manivannan et al.(2015) and Roowi et al.(2010) reported that the morphogenesis and external morphological characteristics of SE were regulated by complex biochemistry and molecules inside the cell tissues. Therefore, it is very important to understand its physiological mechanism in the regulation of this process.

Sugar is an indispensable energy substance and carbon skeleton for embryonic development (Bartos et al., 2018), and it can regulate cell osmotic pressure (Iraqi and Tremblay, 2001), protect cell membrane integrity (Bartos et al., 2018), and determine the shape and developmental fate of cell (Baluska et al., 2003). Proteins affect embryo development (Cangahuala-Inocente et al., 2014), embryo morphogenesis (Cangahuala-Inocente et al., 2014), embryo maturation (Jiménez, 2001) and cell signal transduction (Cangahuala-Inocente et al., 2014). Previous studies have confirmed that the changes in their contents could be used as a marker of different SE stages (Jiménez, 2001; Cangahuala-Inocente et al., 2014).

ROS, such as • OH, H_2_O_2_, and O_2_- continuously occur in mitochondria, chloroplasts and peroxisomes of plant cells as byproducts of various metabolic pathways of respiration and photosynthesis (Libik et al., 2005; Zhang et al., 2010). H_2_O_2_ plays the role of a secondary messenger in cell signaling transduction, affects gene expression (Libik et al., 2005) and stimulates embryonic cell formation (Cui et al., 1999). However, in vitro culture of explants provides a stressful environment, which can stimulate the overproduction of ROS (Fatima et al., 2011), and interfere with intracellular redox homeostasis, normal physiological activities and cell metabolism (Cui et al., 1999). Plants have an antioxidant system to remove excess ROS to maintain the balance of ROS contents in cells (Manivannan et al., 2015). In recent years, changes in the activities of SOD, CAT, POD and APX had been shown to be highly correlated with the ROS elimination at different SE stages (Cui et al., 1999; Libik et al., 2005).

Exogenous hormones play an irreplaceable role in SE induction, and their levels affect the balance of endogenous hormones in plants, regulate plant growth and morphogenesis (Guo et al., 2017), and even determine the developmental fate of cells (Grzyb et al., 2018). Auxin triggers SE induction (Nic-Can and Loyola-Vargas, 2016), and its asymmetric distribution plays key roles in the polar transport of auxin, stem cell formation (Su and Zhang, 2009) and SE induction (Choi et al., 2001). Like auxin, cytokinin is a key regulatory factor for SE (Jiménez, 2005), and the balance between cytokinin and auxin determines the dedifferentiation and redifferentiation states of cells. In addition, ABA and GAs play a positive role in SE maturation and germination (Zhang et al., 2010; Kim et al., 2019). Although endogenous hormones have made some achievements in SE induction and regulation, their regulatory mechanism is still not well understood.

Biochemical events during SE in *O. henryi* were determined to gain a better understanding of the role of the energy substance, ROS, enzyme activities and endogenous hormones in triggering the embryogenic pathway, and they could serve as markers in the determination of different stages of somatic embryo development. Moreover, the histochemical changes in the accumulation and distribution of starch and protein granules were examined at different SE stages, and the changes in starch and protein contents were assessed at different SE stages based on the quantitative and qualitative analysis.

## RESULTS

### Energy substance

The contents of soluble sugar, starch and protein were significantly different at different SE stages (EC -GE -CE) (Fig. 2). The contents of soluble sugar and starch decreased gradually with SE development, and the soluble protein contents increased, which indicated that carbohydrate substances were consumed by somatic embryo growth and development while the increase in protein content offered material conditions for the maturation and germination of somatic embryos. Compared with EC, the contents of soluble sugar, starch and protein were lower in NEC, which indicated that energy substances were the material basis for EC transformation to the somatic embryo; in contrast, the contents of soluble sugar, starch and protein were significantly higher in SC than EC, indicating that the high-energy substance contents facilitated SC formation; however, the soluble protein content was lowest in BC, which indicated that BC activity was low, and might be a cause of its final necrosis.

**Fig. 1.**
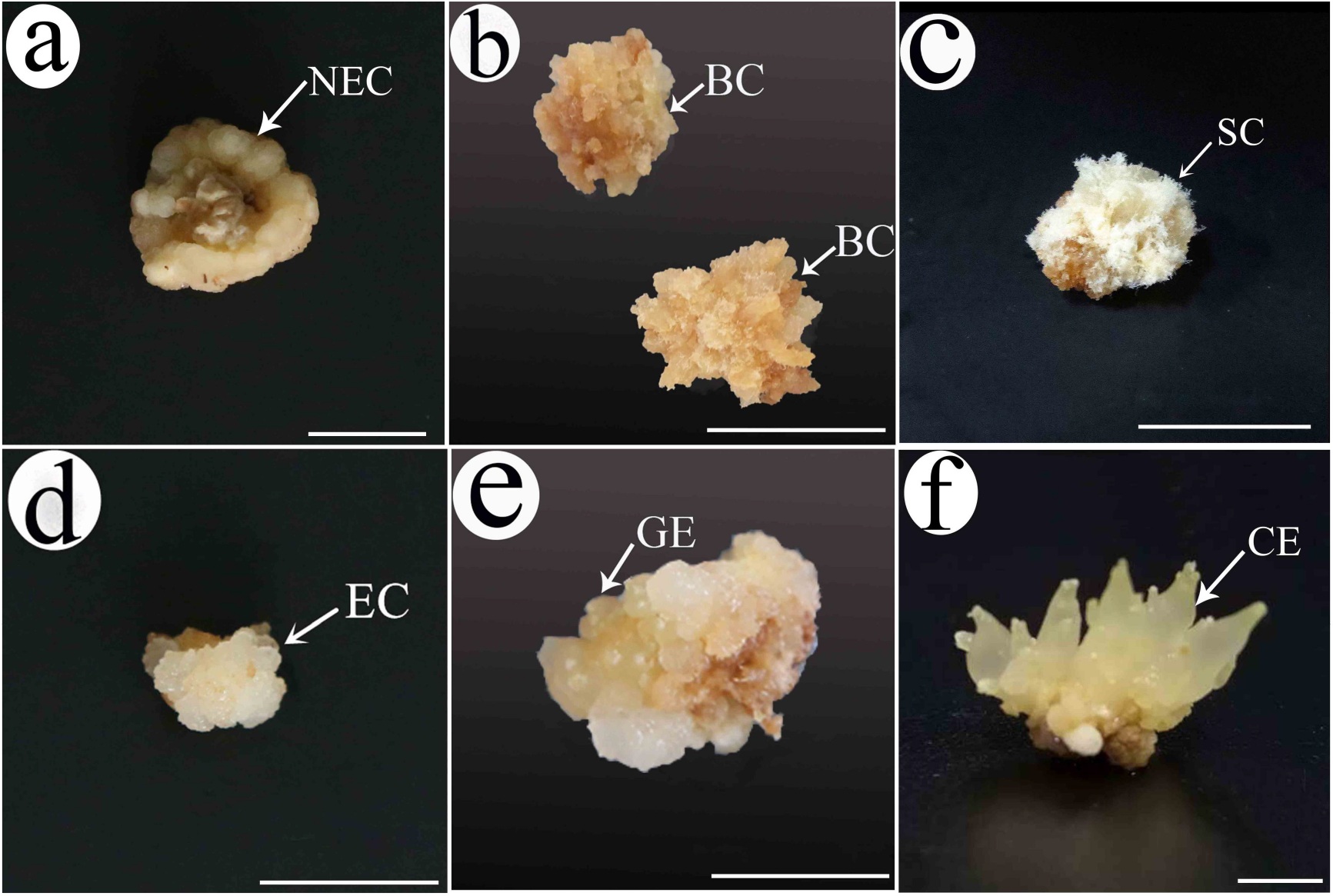
Somatic embryogenesis in *O. henryi*. (a-**d**) The mature embryos were inoculated into B5 medium supplemented with 30 g/L sucrose, 500 mg/L glutamine, 2.5 g/L phytagel, 0.2 mg/L 6-benzylaminopurine (BA) and 2.0 mg/L 2,4-dichlorophenoxyacetic acid (2,4-D) for callus induction. After 6 weeks, callus were transferred into B5 medium supplemented with 0.5 mg/L kinetin (KT), 1.0 mg/L 2,4-D for proliferation induction. After 4 weeks, non-embryogenic callus **(a)**, browning callus **(b)**, snowflake callus **(c)**, and embryogenic callus **(d)** were obtained. **(e)** Globular embryo was obtained after embryogenic callus continued to be cultured in B5 medium supplemented with 0.5 mg/L kinetin (KT), 1.0 mg/L 2,4-D for 6 weeks in the dark at 25 ± 2°C. **(f)** Cotyledon embryo was obtained after globular embryo was cultured in B5 differentiation medium supplemented with 0.5 mg/L thidiazuron (TDZ) and 0.2 mg/L naphthaleneacetic acid (NAA) for 60 days.

**Fig. 2.**
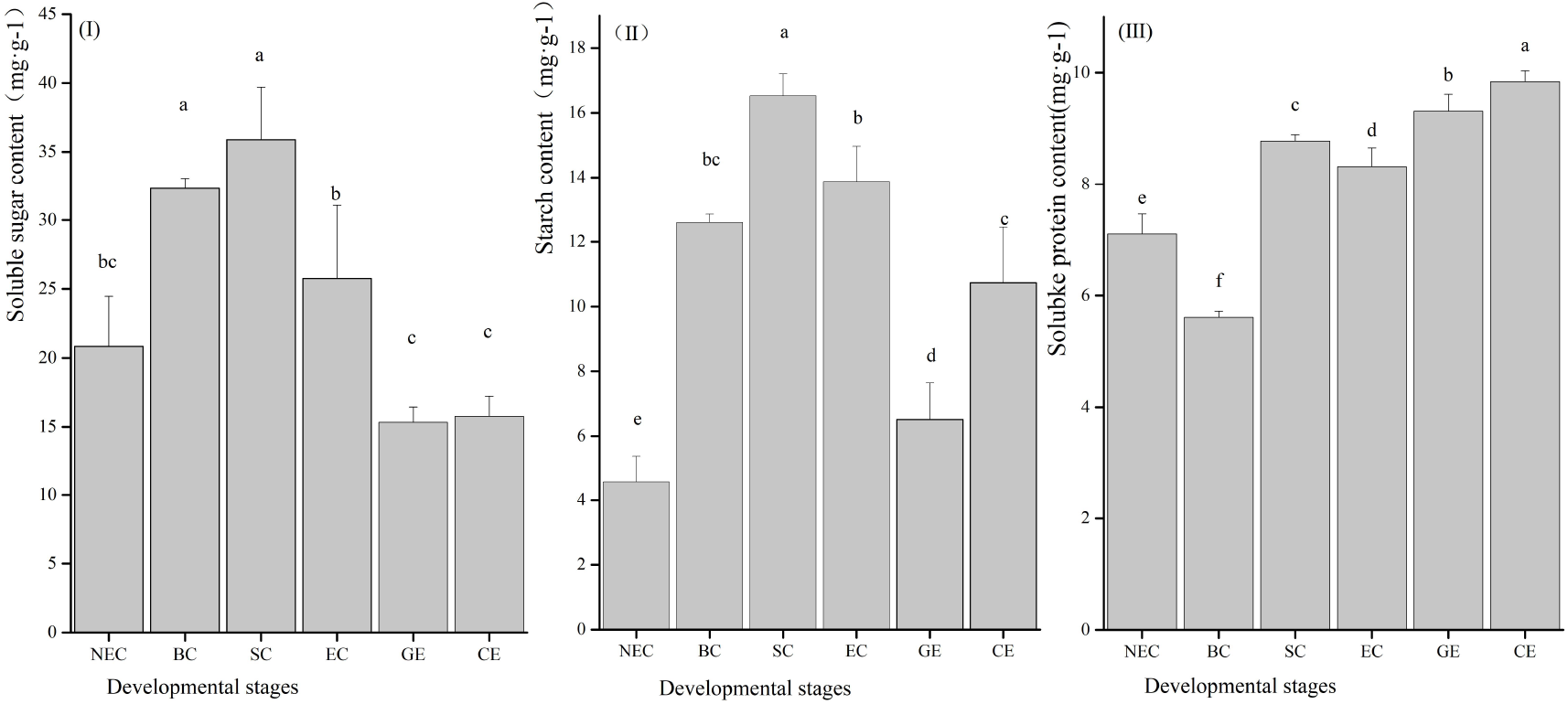
Contents of soluble sugar, starch and protein during somatic embryogenesis in *O. henryi*. **(I)** Soluble sugar contents at different SE stages, such as EC, GE, and CE, and in abnormal tissues, such as NEC, BC, and SC. Bars represent the mean±standard error of three independent experiments, bars denoted by the same letter are not significantly different using the Tukey’s test (p< 0.05). **(II)** Starch contents at different SE stages, such as EC, GE, and CE, and in abnormal tissues, such as NEC, BC, and SC. Bars represent the mean±standard error of three independent experiments, bars denoted by the same letter are not significantly different using the Tukey’s test (p< 0.05). **(III)** Protein contents at different SE stages, such as EC, GE, and CE, and in abnormal tissues, such as NEC, BC, and SC. Bars represent the mean±standard error of three independent experiments, bars denoted by the same letter are not significantly different using the Tukey’s test (p< 0.05).

### H_2_O_2_ contents and enzyme activities

The H_2_O_2_ content and activities of PPO, SOD, APX, POD, and CAT were significantly different at different SE stages (Fig. 3). H_2_O_2_ content increased gradually, whereas the activities of PPO, SOD, APX and POD decreased gradually with SE development. CAT activity, however, was the highest in GE. It is worth noting that APX activity decreased sharply in GE and CE, which was 15% and 23% of EC, respectively. There were significant differences in APX and CAT activities between EC and GE. Among them, APX activity in EC was 570% higher than in GE, and CAT activity was 89% higher in GE than EC.

**Fig. 3.**
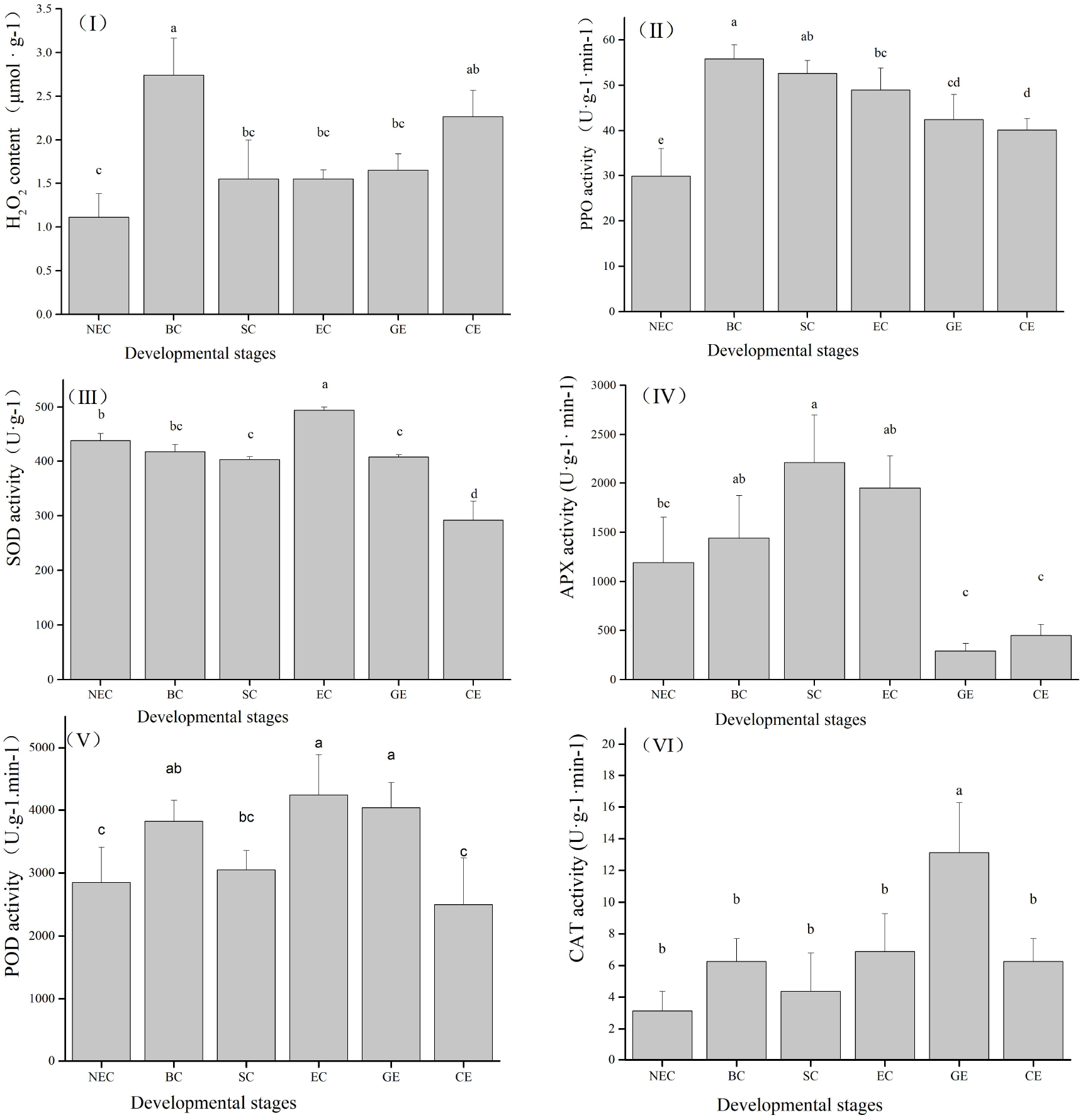
H_2_O_2_ content and enzymatic activity during somatic embryogenesis in *O. henryi*. **(I)** H_2_O_2_ contents at different SE stages, such as EC, GE, and CE, and in abnormal tissues, such as NEC, BC, and SC. Bars represent the mean ± standard error of three independent experiments, bars denoted by the same letter are not significantly different using the Tukey’s test (p< 0.05). **(II)** Polyphenoloxidase activity at different SE stages, such as EC, GE, and CE and in abnormal tissues, such as NEC, BC, and SC. Bars represent the mean ± standard error of four independent experiments, bars denoted by the same letter are not significantly different using the Tukey’s test (p< 0.05). **(III)** Superoxide dismutase activity at different SE stages, such as EC, GE, and CE and in abnormal tissues, such as NEC, BC, and SC. Bars represent the mean ± standard error, bars denoted by the same letter are not significantly different using the Tukey’s test (p< 0.05). **(IV)** Ascorbate peroxidase activity at different SE stages, such as EC, GE, and CE and in abnormal tissues, such as NEC, BC, and SC. Bars represent the mean ± standard error of four independent experiments, bars denoted by the same letter are not significantly different using the Tukey’s test (p< 0.05). **(V)** POD activity at different SE stages, such as EC, GE, and CE and in abnormal tissues, such as NEC, BC, and SC. Bars represent the mean ± standard error of three independent experiments, bars denoted by the same letter are not significantly different using the Tukey’s test (p< 0.05). **(VI)** CAT activity at different SE stages, such as EC, GE, and CE and in abnormal tissues, such as NEC, BC, and SC. Bars represent the mean ± standard error of four independent experiments, bars denoted by the same letter are not significantly different using the Tukey’s test (p< 0.05).

An increase in contents by 36% (H_2_O_2_) and activities by 63% (PPO), 13% (SOD), 64% (APX), 48% (POD), and 115% (CAT) were noticed in EC compared with NEC, which indicated that the high H_2_O_2_ contents and the high enzyme activities could promote SE. The H_2_O_2_ content and PPO activity were highest in BC, which might be the fundamental reason for callus turning into BC. Compared with EC, SOD and POD activities of SC were significantly decreased, which indicated that the low activities of SOD and POD affected SC formation.

### Endogenous hormones

The contents of AUX, CKs, GAs and ABA were significantly different at different SE stages (Fig. 4). Their total contents showed an increasing trend. Except for AUX total contents, which were lowest in GE, the total contents of all endogenous hormones were lowest in EC and highest in CE. These findings indicated that a relatively high AUX content and low contents of CKs, GAs and ABA were conducive to EC induction, a high AUX content was the premise for somatic embryo development and polarity establishment, a high CKs content regulated the differentiation of somatic embryos in the latter stage, and high contents of GAs and ABA promoted the maturation and germination of somatic embryos in the latter stage.

**Fig. 4.**
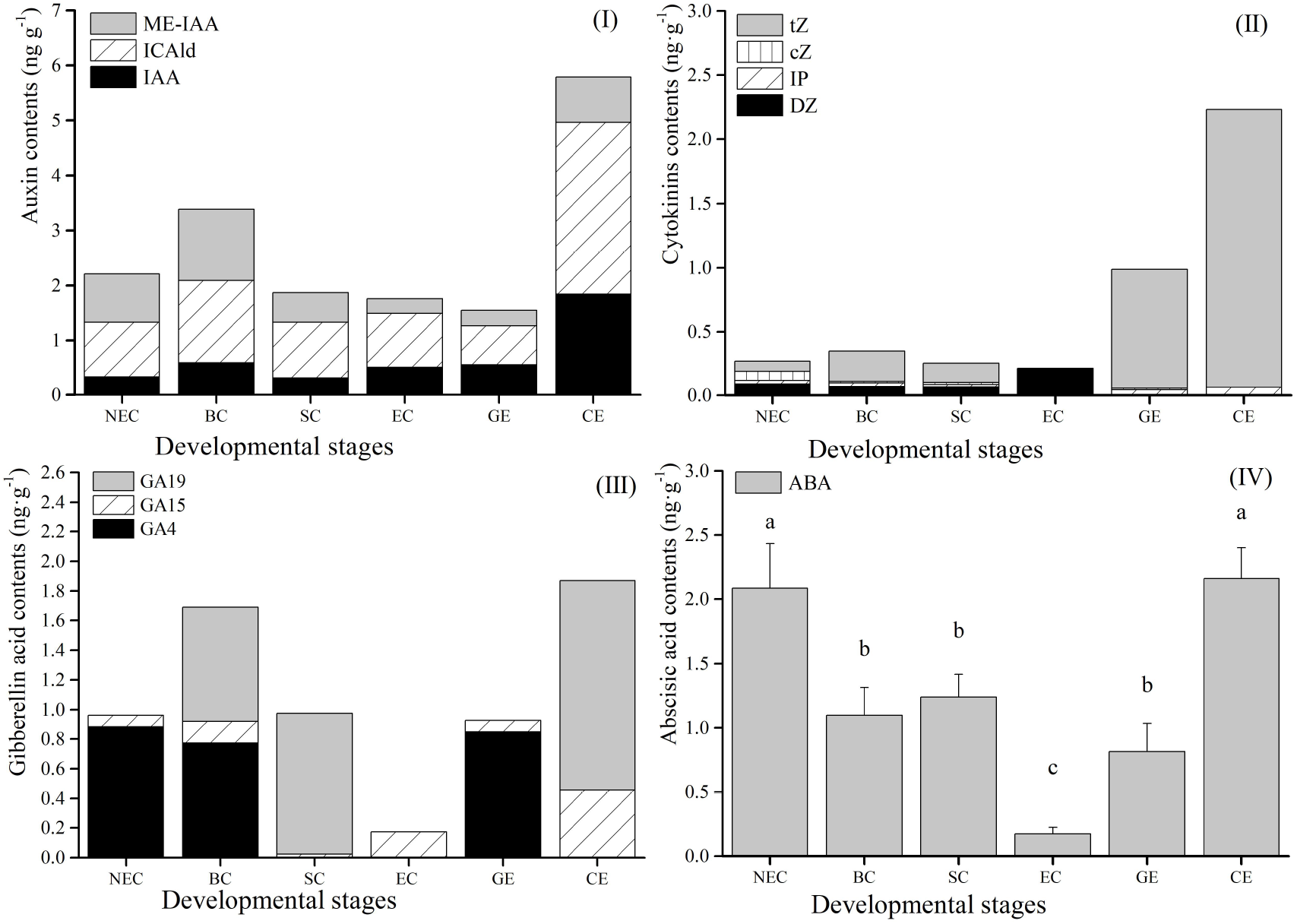
Endogenous hormones contents during somatic embryogenesis in *O. henryi*. **(I)** Auxin (IAA, ICAld and ME-IAA) contents at different SE stages, such as EC, GE, and CE, and in abnormal tissues, such as NEC, BC, and SC. **(II)** Cytokinin (cZ, tZ, DZ and IP) contents at different SE stages, such as EC, GE, and CE, and in abnormal tissues, such as NEC, BC, and SC. **(III)** Gibberellin acid (GA4, GA15 and GA19) contents at different SE stages, such as EC, GE, and CE, and in abnormal tissues, such as NEC, BC, and SC. **(IV)** Abscisic acid (ABA) contents at different SE stages, such as EC, GE, and CE, and in abnormal tissues, such as NEC, BC, and SC. Data are the means ± standard error of three replicates. The use of the same lowercase letters indicates that the values were not significantly different according to Tukey’s test (p < 0.05).

Compared with EC, the contents of AUX and CKs were similar in NEC and SC, and IAA contents in AUX and DZ contents in CKs were higher in EC; the contents of AUX and CKs in BC were significantly higher than those in EC. In addition, the contents of ME-IAA and tZ were lower in EC than NEC, BC and SC. These results showed that NEC formation was related to the low IAA content; the callus with high contents of AUX and CKs was not conducive to EC formation; while the callus with low contents of IAA and DZ, and high tZ content could easily form SC. Moreover, the contents of GAs and ABA in NEC, BC and SC were more than 6 times that of EC, which indicated that the high contents of GAs and ABA inhibited EC induction.

### Endogenous hormones ratios

The interaction between endogenous hormones can be expressed by their ratios to evaluate the balance of endogenous hormones at different SE stages. The ratios of IAA/ABA, IAA/GAs, AUX/GAs and AUX/ABA decreased gradually. The ratios of IAA/CKs, AUX/CKs, ABA/CKs and GAs/CKs first increased and then decreased. The ratios of IAA/CKs and AUX/CKs were the lowest in CE, and the ratios of ABA/CKs, GAs/CKs were lowest in EC. These results indicated that the high ratios of IAA/ABA, IAA/GAs, AUX/GAs, and AUX/ABA and low ratios of ABA/CKs and GAs/CKs were conducive to EC induction at an early stage of SE. The low ratios of IAA/CKs and AUX/CKs promoted maturation and germination of the somatic embryo.

Compared with EC, the ratios of IAA/ABA, IAA/GAs, AUX/GAs, and AUX/ABA decreased significantly in NEC, BC, and SC, and the ratios of ABA/CKs and GAs/CKs were higher. In addition, the ABA/GAs ratio was higher in NEC and lower in BC. These fundings indicated that the low ratios of IAA/ABA, IAA/GAs, AUX/GAs, and AUX/ABA and the high ratios of ABA/CKs and GAs/CKs were responsible for the inability of the callus to form EC; however, an ABA/GAs ratio that was too high or too low ABA/GAs ratio was not conducive to SE induction.

### Histochemical studies

#### Starch granules

As shown in Fig.5, starch granules were mainly distributed in the cell wall and intercellular space, followed by an uneven distribution in cytoplasm and reduced levels in the nucleus. These findings showed that carbohydrates were the main components of the carbon skeleton, and cytoplasm was the main site of carbohydrate synthesis.

**Fig. 5.**
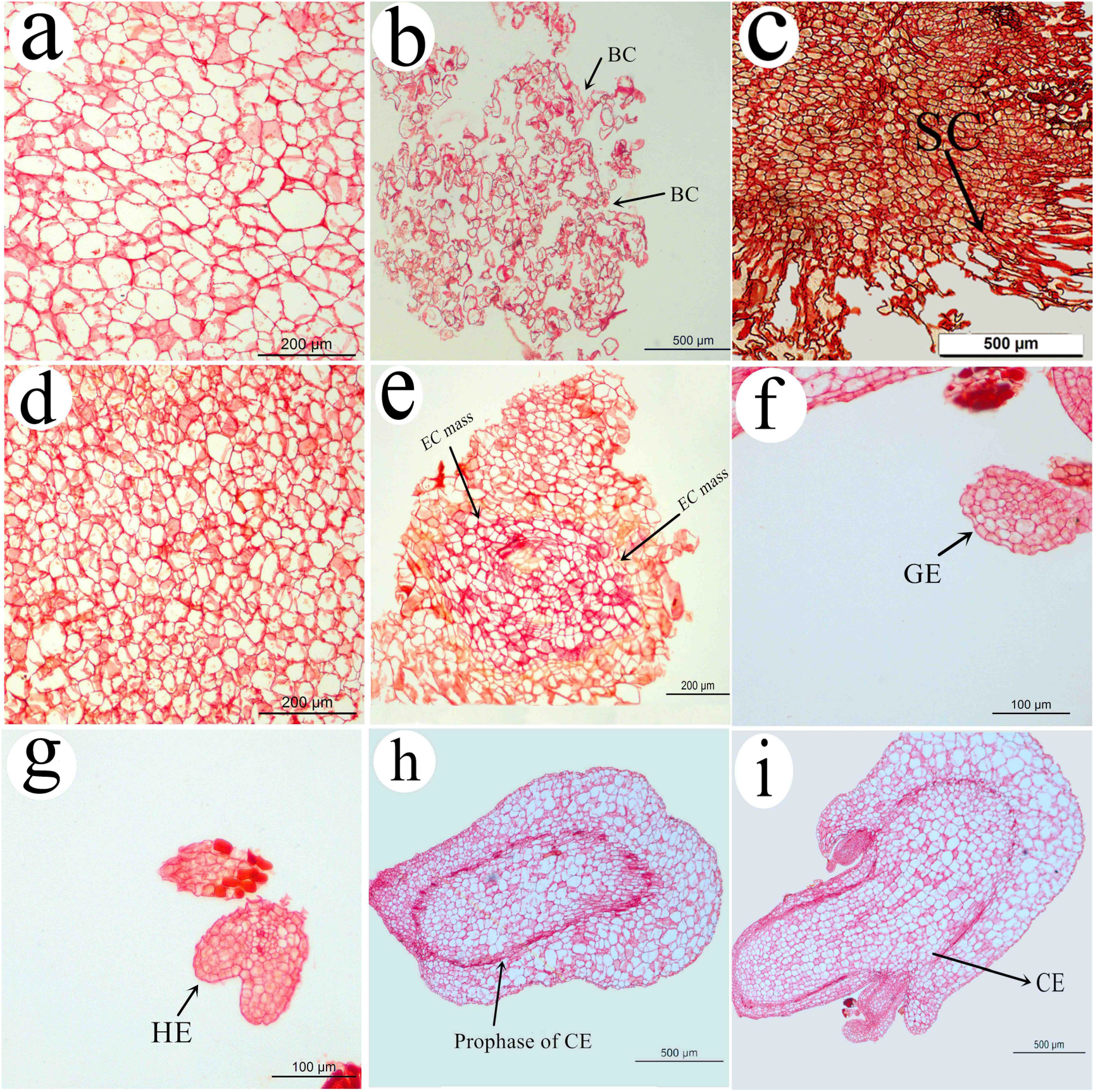
Starch grains staining during somatic embryogenesis in *O. henryi*. Starch granules were mainly distributed in cell wall and intercellular space, followed by uneven distribution in cytoplasm, less in nucleus, and then the meristem staining was dark. **(a)** Non-embryogenic callus. **(b)** Embryogenic callus. **(c)** Embryogenic callus. **(d)** Browning callus. **(e)** Snowflake callus. **(f)** Globular embryo. **(g)** Heart-shaped embryo. **(h)** Prophase of CE. **(i)** Cotyledon embryo.

At different SE stages, the starch granules staining was dark in the EC, and starch granule staining of EC masses (Fig. 5-c) was especially darker than in the surrounding tissues, which indicated that EC masses stored sufficient nutrients for the growth and development of SE. The starch granule staining gradually lightened with SE development, but the meristem staining (Fig. 5f-i) was dark, which confirmed that the meristem was active and might provide the best explanation for the strong embryogenic response of the shoot apical meristem (SAM).

Compared with EC, the starch granule staining was lightest in BC, followed by NEC, but it was darker in SC. These findings indicated that the abundant energy substances were the basis for callus formation of EC, but the callus with excessive energy substances could easily form SC.

#### Protein granules

As shown in Fig.6, the protein granule staining was relatively light, which was most obvious in the nucleus, followed by the cytoplasm, which indicated that proteins in the nucleus were most abundant. This feature was related to the need for a large number of proteins and enzymes for DNA replication and transcription in the nucleus.

**Fig. 6.**
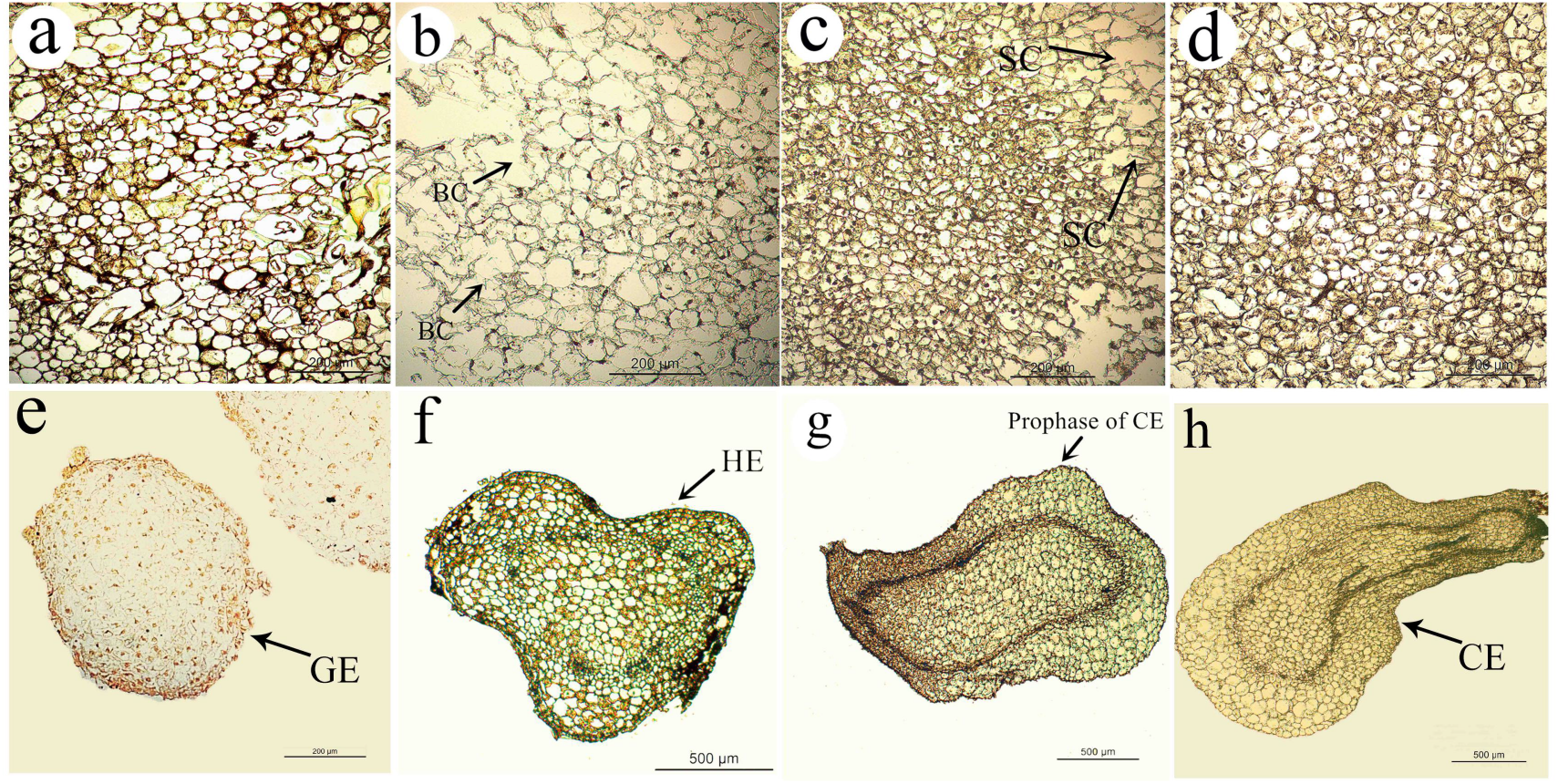
Protein grains staining during somatic embryogenesis in *O. henryi*. Protein granules staining was relatively light, which was the most obvious in nucleus, followed by cytoplasm, and then there were more protein grains accumulated in the meristems. **(a)** Non-embryogenic callus. **(b)** Embryogenic callus. **(c)** Browning callus. **(d)** Snowflake callus. **(e)** Globular embryo. **(f)** Heart-shaped embryo. **(g)** Prophase of cotyledon embryo. **(h)** Cotyledon embryo.

With SE development, the protein grain staining became gradually darkened (Fig. 6d-h), with the darkest staining observed in CE. More protein grains accumulated in the meristems, which indicated their strong activity and vigorous metabolism.

Compared with EC, the protein granules staining were light in NEC and BC, and with greater numbers of vacuolar cells, the numbers of nuclei in the cell were reduced; however, it was darker in SC. These results indicated that the callus with the low protein content and activity affected the ability of the callus to form EC, but the callus with the high protein content could easily form SC.

## DISCUSSION

### Energy substances

C and N participate in the metabolic energy, carbon skeletons and biochemical signaling during plant growth and development (Bartos et al., 2018), and they are closely related to the growth and development of SE and morphogenesis (Cangahuala-Inocente et al., 2014). In this study, the contents of soluble sugar and starch gradually decreased with SE development, similarly to findings reported in *Acca sellowiana* (Cangahuala-Inocente et al., 2014) and spruce (Hazubska-Przybyl et al., 2016). This decrease was related to the consumption of a large amount of energy for the growth and development of SE. However, the protein content increased with SE development, similarly to findings reported in *Eucalyptus globulus* (Pinto et al., 2010), *Coffea arabica* (Bartos et al., 2018), *Catharanthus roseus* (Fatima et al., 2011) and *Cordyline australis* (Warchoł et al., 2015). These phenomena could be related to the synthesis of late embryogenesis abundant protein (LEA) and reserve proteins, which might play an important role in embryonic development and protect embryos against dehydration (Gomes et al., 2014); however, the protein content in late stages of SE were decreased in *Acca sellowiana* (Cangahuala-Inocente et al., 2014) and *Persian walnut* (Jariteh et al., 2015).

EC induction was the basis of SE. The contents of soluble sugar, starch and protein were higher in EC than NEC, similarly to findings reported in banana (Wang et al., 2013), alfalfa (Martin et al., 2000) and *Catharanthus roseus* (Fatima et al., 2011), because it was difficult or impossible for NEC to develop into the somatic embryo. These findings showed that the accumulation of energy substances was the material basis for EC transformation into the somatic embryo. Compared with EC, the contents of starch and protein were lower in BC, which indicated that BC activity was low and thus affected the synthesis of starch and protein, which may explain the final necrosis of BC. In contrast, the contents of soluble sugar, starch and protein increased significantly in SC, but SC could not further develop into somatic embryos, which confirmed that the accumulation of energy substances affected the cell activity and morphogenesis. These results indicated that too low or too high contents of energy substances were not conducive to the further growth of callus. A previous study confirmed that the sucrose concentration affected carbon metabolism in EC, and medium supplemented with different sucrose concentrations was needed to reduce the frequency of SC, although the frequency of SC (9%) was significantly reduced at a low sucrose concentration (20g/L), the SE induction rate (19%) also reduced.

### H_2_O_2_ contents and enzyme activities

The H_2_O_2_ content increased in the late stage of SE, and similar results were reported in wolfberry (Cui et al., 1999) and Begonia (Manivannan et al., 2015), which indicated that ROS metabolism played a decisive role in cell differentiation and development (Cui et al., 1999). The activities of SOD and APX decreased in GE and CE. Fatima et al.(2011) reported similar results in *Catharanthus roseus*; APX activity significantly decreased, which might be related to the high activities of POD and CAT in GE and CE, which consumed excessive H_2_O_2_, leading to the decrease in APX activity. A similar phenomenon was reported in *Citrus reticulata* (Liu, 2016).

H_2_O_2_ content and the activities of PPO, SOD, APX, POD and CAT in the EC of *O. henryi* were higher than those in NEC, which indicated that the callus with the low H_2_O_2_ content and low enzyme activities were not sufficient to induce SE. The H_2_O_2_ accumulation could trigger the stress response, play a role as a secondary messenger involved in cell signaling and promote SE development. Similar results have been reported in *Albizia lebbeck* (Saeed and Shahzad, 2015) and *Mesembryanthemum crystallinum* (Libik et al., 2005). Compared with EC, higher H_2_O_2_ contents, higher PPO activity and lower antioxidant enzyme activities were detected in BC, which caused BC to be in a state of high oxidative stress. Wichers et al.(1985) reported that PPO activity increased in cell culture of *Mucuna pruriens* during cell senescence, which might represent the main reasons for callus transformation into BC and even necrosis. To reduce the BC frequency, medium supplemented with anti-browning agents, such as polyvinylpyrrolidone, ascorbic acid and activated carbon, reduced the frequency of callus transformation into BC, but the results were not significant differences (data not shown). Compared with EC, SOD and POD activities decreased significantly in SC, and therefore, we speculated that callus with low SOD and POD activities could easily form SC. Fatima et al.(2011) reported that POD was a marker involved in cell morphogenesis.

### Endogenous hormones

Endogenous hormone levels regulate the dedifferentiation, redifferentiation and embryogenesis of somatic cells at different SE stages (Grzyb et al., 2018). At different SE stages of *O. henryi*, IAA content increased significantly in CE, which was consistent with the results of SE maturation of *Norway Spruce* (Vondrakova et al., 2018) and larch (Aderkas et al., 2001), indicating that auxin is related to the establishment of embryo polarity (Vondrakova et al., 2018). Auxin is required for SE maturation to mediate SAM by establishing the homeodomain transcription factor WUSCHEL (WUS), which is essential for stem cell initiation and meristem maintenance (Lee and Clark, 2015). Similarly, CKs contents increased significantly in CE, which is consistent with a report on SE maturation and germination in spruce (Vondrakova et al., 2018). Recent advances have shown that CKs are essential factors that regulate shoot initiation via WUS expression (Cheng et al., 2013), and the CKs response factors ARABIDOPSIS RESPONSE REGULATORs (ARRs) directly activate WUS expression and thus initiate SAM formation (Zhang et al., 2017; Liu et al., 2018). Some studies had shown that high GAs levels can inhibit SE induction, but GAs have a positive effect on the maturation and germination of embryos (Kim et al., 2019), as also confirmed in our study. ABA content was also the highest in CE, indicating that ABA could promote the maturation and differentiation of embryos. Vondrakova et al.(2018) found that ABA content was highest in the SE maturation stage of spruce. Cheng (2016) reported that the ABA content increased significantly after EC formation in cotton, and the genes related to ABA synthesis and signal transduction were also differentially expressed in cotton. The ABA2, NCED, PP2C, ABF, PYR/PYL and ABA receptor family were upregulated in the embryo.

We found that the IAA contentwas higher in EC than NEC, which was consistent with the results obtained for *Cyathea delgadii* (Grzyb et al., 2017). Su and Zhang(2009) reported that auxin gradients must be established for EC induction, and expressive of WUS gene was correlated with auxin gradients and initiated the polar localization of PIN1in EC. These findings indicated that endogenous auxin had important significance for WUS gene and SE induction. The contents of tZ and cZ in NEC were higher than in EC, similarly to findings reported in *Prunus persica* (Pérez-Jiménez et al., 2013), this result was also supported by the research of Fraga et al.(2016), who found that the decrease in the Z level was related to the embryogenic potential. Compared with EC, BC with high contents of AUX and CKs was not conducive to EC formation; while the callus with low contents of IAA and DZ, and high tZ content could easily form SC. In addition, the contents of GAs and ABA in NEC, BC and SC were more than 6 times that of EC, which indicated that the high contents of GAs and ABA inhibited EC induction. Similar results had shown that the higher ABA content in NEC was also found in *Hevea brasiliensis* (Etienne et al., 1993) and *Cyathea delgadii* (Grzyb et al., 2017), while Pérez-Jiménez et al.(2013) found no significant difference in ABA content between NEC and EC in *Prunus persica*. In contrast, Nakagawa et al.(2001) found a higher ABA level in EC in *Cucumis melo*. These difference may depend on the sensitivity of different species to ABA and the difference in time and space. The content difference of these endogenous hormones in different SE stages and different callus can provide a reference for SE induction and the application of exogenous hormone.

### Endogenous hormone ratios

The ratios of endogenous hormones contents can be used as a physiological index to regulate the dedifferentiation and redifferentiation of the cell (Grzyb et al., 2018), and evaluate embryogenesis (Pérez-Jiménez et al., 2013). At different SE stages of *O. henryi*, the high ratios of IAA/ABA, IAA/GAs, AUX/GAs, and AUX/ABA and low ratios of ABA/CKs and GAs/CKs were favorable for EC induction; however, the ratios of IAA/CKs and AUX/CKs were lower in CE and promoted SE maturation and differentiation, which indicated that low AUX and high CKs contents were needed for SE maturation and differentiation, consistent with the actual theory.

The IAA/ABA ratio was significantly higher in EC than NEC, and similar results were found in *Corylus avellana* (Centeno et al., 1997); In contrast, the ABA/CKs ratio was significantly higher in NEC than EC, and similar results have been reported by Grzyb et al. (2017). Guo et al.(2018) reported that the ABA/GAs ratio can reflect the growth and dormancy of plant, as well as determine the developmental pattern of the embryo or late stages of the embryo (Zheng et al., 2013). Our study showed that the callus with low ratios of IAA/ABA, IAA/GAs, AUX/GAs, and AUX/ABA, and high ratios of ABA/CKs and GAs/CKs could not form EC, and an ABA/GAs ratio that was too high or too low was not conducive to SE induction.

### Histochemical studies

Starch and protein are active metabolites in cell growth and morphogenesis. The starch granule staining gradually lightened with SE development of *O. henryi*, and the protein grain staining gradually darkened. Similar findings have been reported in *Picea sitchensis* SE (Liu, 2009). The changes at different SE stages can be used as a biochemical marker in *O. henryi* and provide important information for SE induction.

Compared with EC, the starch and protein granules staining were lighter in NEC. Similar results have been reported in *Picea sitchensis* (Liu, 2009) and *Albizia lebbeck* (Saeed and Shahzad, 2015); similarly, the starch and protein granules staining were lighter in BC and darker in SC. These phenomena were consistent with the results of our quantitative analysis, which indicated that contents of energy substances that were too low or too high were not conducive to callus development.

In summary, SE was a complex physiological process, and a single or two indexes was insufficient to evaluate the SE mechanism in *O. henryi*. The physiological indexes interacted with each other, and the results showed that energy substances were the material basis of SE, which affected the enzyme activities, regulated ROS metabolism, and thus regulated the morphogenesis and development of somatic embryos. In addition, the contents and ratios of endogenous hormones affected the dedifferentiation, dedifferentiation and embryogenesis of somatic cells. The formation of NEC was related to low contents of energy substances. H_2_O_2_ and low enzyme activities; the high contents of soluble sugar, H_2_O_2_, AUX, CKs, and high PPO activity and low content of soluble protein were the basic causes of BC formation; the high-energy substance contents and low activities of SOD and POD could easily form SC. The high contents of Gas and ABA, high ratios of ABA/CKs and GAs/CKs, and low ratios of IAA/ABA, IAA/GAs, AUX/GAs, and AUX/ABA were explanations for the inability of the callus to form EC. Based on the above conclusion, the EC induction rate and frequency of EC transformation into a somatic embryo in *O. henryi* might be regulated and improved by adjusting the external conditions, such as the medium composition and culture environment, especially the application of exogenous hormone types and concentrations.

## MATERIALS AND METHODS

### Plant materials

Mature seeds were collected from a 120-150-year-old *O. henryi* tree growing in Mengguan town, Guiyang city (latitude: 26°14’23”N, longitude: 106°25’12’’W) on Nov 2017. The mature seeds were treated with concentrated H_2_S0_4_ for 1 h, washed with tap water for 30 min, and disinfected in 75% alcohol for 1 min and then in 2% NaClO for 8 min, followed by five rinses in sterile distilled water. Because the seed coat is hard, the mature seeds were soaked in sterile water for 24 h to make them swell and soften, which was conducive to stripping the mature embryos. Mature embryos were used as the explant for SE induction (Wu et al., 2020).

The mature embryos were inoculated into B5 medium (Gamborg et al., 1968) supplemented with 30 g/L sucrose, 500 mg/L glutamine, 2.5 g/L phytagel, 0.2 mg/L 6-benzylaminopurine (BA) and 2.0 mg/L 2,4-dichlorophenoxyacetic acid (2,4-D) for callus induction. The pH values of the medium were adjusted to 5.90±0.1 before autoclaving 121°C for 20 min. After 6 weeks, callus were transferred into B5 medium supplemented with 0.5 mg/L kinetin (KT), 1.0 mg/L 2,4-D for callus proliferation induction; after 4 weeks, NEC (Fig. 1-a), BC (Fig. 1-b), SC (Fig. 1-c) and EC (Fig. 1-d) were collected. GE (Fig. 1-e) was obtained after EC continued to be cultured in the above medium for 2 weeks in the dark at 25 ± 2°C. CE (Fig. 1-f) was obtained after GE was cultured in B5 differentiation medium supplemented with 0.5 mg/L thidiazuron (TDZ) and 0.2 mg/L naphthaleneacetic acid (NAA) for 60 days.

The different tissues were collected, the culture medium of tissue surface was cleaned with deionized water. The tissues were placed in 5 ml centrifuge tubes, quickly frozen in liquid nitrogen, and then transferred to −80 °C freezer for storage.

### Biochemical analyses

The contents of energy substances, ROS, endogenous hormones and enzyme activities in different tissues, such as NEC, BC, SC, EC, GE and CE, were determined. Fresh tissues of 0.2-0.5g were obtained to detect and analyze biochemical events, with 3-5 replications performed per treatment.

### Energy substances

The contents of soluble sugar and starch were determined using the sulfuric acid-anthrone method (Liu et al., 2016), and the soluble protein content was determined using Coomassie brilliant blue (Bradford, 1976).

### H_2_O_2_ contents and enzyme activities

H_2_O_2_ was determined according to the method of Ferguson et al. (1983) method. SOD activity was determined using the nitroblue tetrazolium method (Dhindsa et al., 1981). PPO activity was determined by pyrocatechol colorimetry (Liu et al., 2016). POD activity was measured using the guaiacol colorimetric method (Türkan et al., 2005). The determination of APX activity was based on the method of Yoshiyuki and Kozi (1981), and CAT activity was estimated according to Aebi (1984).

### Endogenous hormones

The contents of endogenous hormones, such as AUX (indole-3-acetic acid [IAA], methyl indole-3-acetate [ME-IAA], 3-indolebutyric acid [IBA], indole-3-carboxylic acid [ICA], indole-3-carboxaldehyde [ICAld]), CKs (N6-isopentenyladenine [IP], trans-zeatin [tZ], cis-zeatin [cZ], dihydrozeatin [DZ]), GAs (gibberellin A1 [GA1], GA3, GA4, GA7, GA9, GA15, GA19, GA20, GA24, GA53) and ABA, were measured using the Metware(http://www.metware.cn/) method in different tissues. The tissues were removed from −80 °C freezer and ground into a powder with a grinder (MM 400, Retsch) (30 Hz, 1 min); then, 50 mg of the powdered tissues was added as an internal standard of the appropriate amount and extracted with methanol: water: formic acid = 15:4:1 (v: v: v). The extracts were concentrated and then redissolved in 100 μl 80% methanol aqueous solution and filtered with 0.22 μm PTFE filter membranes. They were then placed in sample bottles and analyzed using liquid chromatography-tandem mass spectrometry (LC-MS/MS). The peak area of each peak represented the relative content of the corresponding hormone. Finally, the qualitative and quantitative analysis results were obtained for all sample hormones.

### Histochemical studies

The different tissues were fixed in FAA (formalin: acetic acid: 50% alcohol, 1:1:18) for 24 h. The tissue was processed with an alcohol-xylol series and embedded in paraffin. Sections (8 μm) were cut on a rotary microtome (Leica, Germany) and dewaxed with xylol. Starch granules were stained with Schiff reagent (Liu, 2009), and protein granules with naphthol yellow S reagent (Liu, 2009). Finally, the sections were mounted with neutral balsam and examined under a DM3000 light microscope (Leica, Germany).

**Table 1.** Ratios of phytohormones during somatic embryogenesis in *O. henryi*. Data are the means (±SD) of three replicates. The use of the same lowercase letters indicates that the values were not significantly different according to Tukey’s test (p < 0.05). CKs represent the sum of cZ, tZ, DZ and IP contents. GAs represent the sum of GA4, GA15 and GA19 contents. AUX represent the sum of IAA, ICAld and ME-IAA contents.

| Phytohormone ratios | NEC | EC | BC | SC | GE | CE |
| --- | --- | --- | --- | --- | --- | --- |
| IAA/ABA | 0.16±0.02b | 3.16±0.58a | 0.57±0.1b | 0.3±0.05b | 0.74±0.2b | 0.86±0.1b |
| IAA/CKs | 1.24±0.09a | 2.38±0.04a | 1.74±0.1a | 1.3±0.3a | 4±2a | 0.88±0.2a |
| IAA/GAs | 0.36±0.04c | 2.94±0.17a | 0.36±0.02c | 0.33±0.03c | 0.6±0.08bc | 1.01±0.13b |
| AUX/CKs | 8.16±0.44a | 8.22±0.26a | 9.89±1.08a | 7.89±1.51a | 10.9±5.26a | 2.72±0.43a |
| AUX/GAs | 2.38±0.62b | 10.15±0.88a | 2±0.06b | 1.93±0.21b | 1.71±0.33b | 3.19±0.76b |
| AUX/ABA | 1.07±0.14b | 10.93±3.52a | 3.16±0.61b | 1.53±0.21b | 2.02±0.63b | 2.7±0.35b |
| ABA/CKs | 7.69±0.68a | 0.81±0.16b | 3.3±0.6ab | 5.2±1.1ab | 5.5±2.7ab | 1±0.1b |
| ABA/GAs | 2.29±0.5a | 0.99±0.16b | 0.65±0.07b | 1.3±0.2ab | 0.9±0.1b | 1.2±0.2ab |
| GAs/CKs | 3.55±0.43a | 0.81±0.04a | 4.93±0.49a | 4.02±0.59a | 6.95±3.33a | 0.88±0.18a |

### Statistical analysis

The results were expressed as the mean ± standard error for 3-4 replications per treatment. They were analyzed by one-way ANOVA followed by Tukey’s test (p < 0.05) with SPSS (version 18).

**Supplemental Data**

**Supplemental Energy substance**

**Supplemental H2O2 contents and enzyme activities**

**Supplemental hormones levels**

## Acknowledgments

We are very grateful to the members of the research group for their help and interesting scientific discussions. This work has benefited from the facilities and expertise of Metware (http://www.metware.cn/) and the experimental condition of the Institute for Forest Resources & Environment of Guizhou.

## Author contributions

Wei X.L conceived the project, designed and supervised the experiments. Wu G.Y., Wang X., Liang X and Wei Y performed the experiments. Wu G.Y and Wang X analyzed data and wrote the manuscript. Wei X.L edited and revised this manuscript.

